# The X chromosome of the German cockroach, Blattella germanica, is homologous to a fly X chromosome despite 400 million years divergence

**DOI:** 10.1101/279737

**Authors:** Richard P Meisel, Pablo J Delclos, Judith R Wexler

## Abstract

**Background:** Sex chromosome evolution is a dynamic process that can proceed at varying rates across lineages. For example, different chromosomes can be sex-linked between closely related species, whereas other sex chromosomes have been conserved for *>*100 million years. Cases of long-term sex chromosome conservation could be informative of factors that constrain sex chromosome evolution. Cytological similarities between the X chromosomes of the German cockroach (*Blattella germanica*) and most flies suggest that they may be homologous—possibly representing an extreme case of long-term conservation.

**Results:** To test the hypothesis that the cockroach and fly X chromosomes are homologous, we analyzed whole genome sequence data from cockroach. We found evidence in both sequencing coverage and heterozygosity that a significant excess of the same genes are on both the cockroach and fly X chromosomes. We also present evidence that the candidate X-linked cockroach genes may be dosage compensated in hemizygous males. Consistent with this hypothesis, three regulators of transcription and chromatin on the fly X chromosome are conserved in the cockroach genome.

**Conclusions:** Our results support our hypothesis that the German cockroach shares the same X chromosome as most flies. This may represent convergent evolution of the X chromosome in the lineages leading to cockroaches and flies. Alternatively, the common ancestor of most insects may have had an X chromosome that resembled the extant cockroach and fly X. Cockroaches and flies diverged ∼400 million years ago, which would be the longest documented conservation of a sex chromosome. Cockroaches and flies have different mechanisms of sex determination, raising the possibility that the X chromosome was conserved despite evolution of the sex determination pathway.

## Background

In species with separate sexes, genetic or environmental cues initiate sexually dimorphic developmental pathways [1, 2]. If the cue is genetic, a sex determining factor may reside on a sex chromosome [3]. For example, in most therian mammals, *SRY* on the Y chromosome initiates the development of the male germline, testes, and secondary sexual traits [4]. In contrast, the dosage of the X chromosome determines the initiation of male or female development in *Drosophila melanogaster* [5–7]. In both taxa, females have the XX genotype, and males are XY. Despite the superficial similarities, the sex chromosomes and genes that initiate the sex determination pathways are not homologous between mammals and *Drosophila* [3]. In addition, some, but not all, animal taxa have evolved mechanisms to compensate for the haploid dose of the X chromosome in males, or Z chromosome in ZW females [8–11].

Sex determining pathways and sex chromosomes can evolve rapidly, often differing between closely related species [2, 3]. Evolutionary transitions in sex determination pathways are often accompanied by corresponding changes in the identity of the sex chromosomes [1, 2, 12]. Transitions in sex determining pathways and turnover of sex chromosomes are well studied across insects, where there is a diversity of sex determination mechanisms [13–16] (**Figure 1**). For example, the genetic factors that initiate sex determination in *Drosophila* do not determine sex in other flies [17– 24]. In addition, the sex chromosomes of *Drosophila* are not homologous to the sex chromosomes of other flies [25–27]. The evolution of a new sex determination mechanism in the lineage leading to *Drosophila* resulted in the transition of the ancestral X chromosome into an autosome, the creation of a new X chromosome from an ancestral autosome, and the evolution of a new mechanism of X chromosome dosage compensation [27, 28].

**Figure 1.**
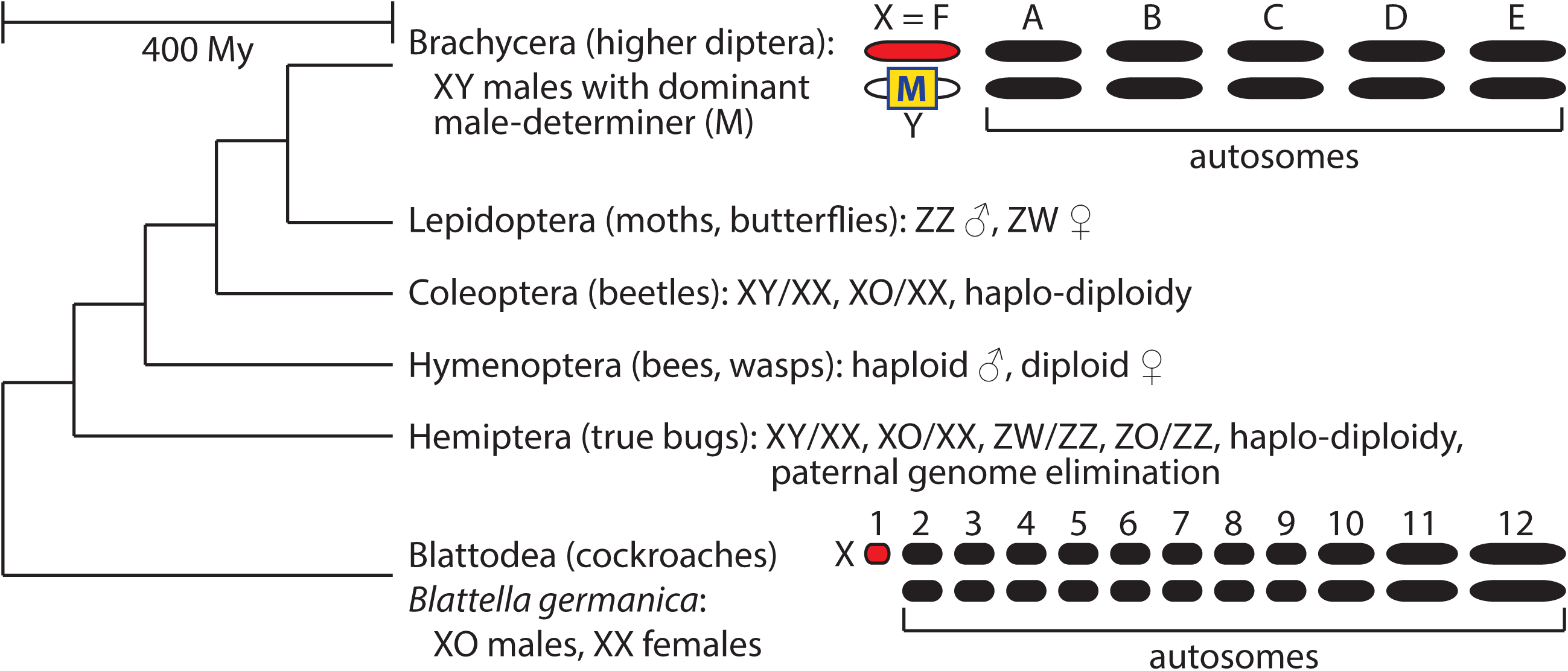
Insect phylogeny and sex chromosomes. Evolutionary relationships and sex chromosome karyotypes of major insect groups. The phylogenetic topology and time to common ancestor are shown [44], but the relative branch lengths are not drawn to scale. Information on insect sex chromosomes and sex determination are reviewed elsewhere [2, 3, 13, 16, 27].

It is most parsimonious to conclude that the ancestral sex determination system of brachyceran dipterans (which includes flies but excludes mosquitoes, craneflies, midges, gnats, etc.) consists of a Y-linked male-determining factor that regulates the splicing of the *transformer* (*tra*) gene product [15, 20, 24, 29–32]. The ancestral male-determining gene of brachyceran flies is yet to be identified, if it is even still present in any extant species. The ancestral brachyceran X chromosome is known as Muller element F [27]. Element F has reverted to an autosome in *D. melanogaster*, where it is also known as chromosome 4 or the “dot” chromosome. The dot chromosome is enriched for heterochromatin and has fewer than 100 genes [33]. Element F is notable because most X chromosomes are gene rich and euchromatic, despite having some differences in gene content from the autosomes [34–36]. This peculiar element F X chromosome has been conserved for *>*150 million years (My) in some fly lineages, but it reverted to an autosome in *Drosophila* when a different chromosome became X-linked [27, 37]. The remainder of the fly genome is organized into five euchromatic chromosomes (or chromosome arms), named Muller elements A–E [38, 39]. Element A is the X chromosome in *D. melanogaster*.

There is some evidence that the X-linked element F is dosage compensated in hemizygous males. In *D. melanogaster*, where element F is autosomal, *Painting of fourth* (*Pof*) encodes an RNA-binding protein that localizes predominantly to element F [40]. *Lucilia cuprina* (Australian sheep blowfly) has the ancestral brachyceran karyotype, with an X-linked element F [41, 42]. Expression of X-linked genes is up-regulated in *L. cuprina* males by the homolog of *Pof* [41, 43]. This dosage compensation is essential for male viability—a loss of function mutation in the *L. cuprina* homolog of *Pof* is male lethal, but viable in females [43].

The German cockroach, *Blattella germanica*, diverged from flies ∼400 My ago (Mya) [44]. Cockroach females are XX and males are XO, i.e., one X and no Y chromosome [13, 45]. This suggests that a dosage-sensitive X-linked factor determines sex in German cockroach, analogous to, but independently evolved from, *Drosophila*. Curiously, the cockroach X chromosome is heterochromatic along most of its length [46], reminiscent of element F, the ancestral brachyceran X chromosome. We tested the hypothesis that the German cockroach X chromosome is homologous to fly element F, which would suggest that a cockroach and most flies share an X chromsomome despite ∼400 My divergence.

## Results

### Decreased sequencing coverage of element F homologs in male cockroaches

We used a differential sequencing coverage approach to identify X chromosome genes in the German cockroach genome assembly. X-linked genes are expected to have half as many male-derived reads mapped to them as female-derived reads because the X chromosome is present in one copy in males and two copies in females [27]. We used available whole genome sequencing data [47] to calculate the relative cover-age of male (*M*) and female (*F*) reads 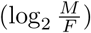 for each annotated cockroach gene (Additional file 1). The mode of the 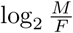 distribution is at 0 (**Figure 2A**), as expected because we recalibrated the 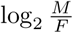 values to have a median of 0 (see Methods). However, there is a heavy shoulder of genes with 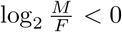, suggesting that X-linked genes are also in the assembly (**Figure 2A**). In total, 3,499 of the 28,141 annotated genes have female-biased coverage 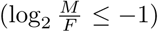, whereas only 1,363 genes have male-biased coverage 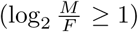, consistent with a heavy shoulder of X-linked genes. Assuming the 1,363 male-biased genes represent the false-positive rate, we expect 2,136/3,499 female-biased genes to be X-linked. This is consistent with the upper-bound of the number of X-linked genes in the cockroach genome— the cockroach X is the smallest of 12 chromosomes [46], which means that fewer than 2,345 genes (28,141/12) should be X-linked.

**Figure 2.**
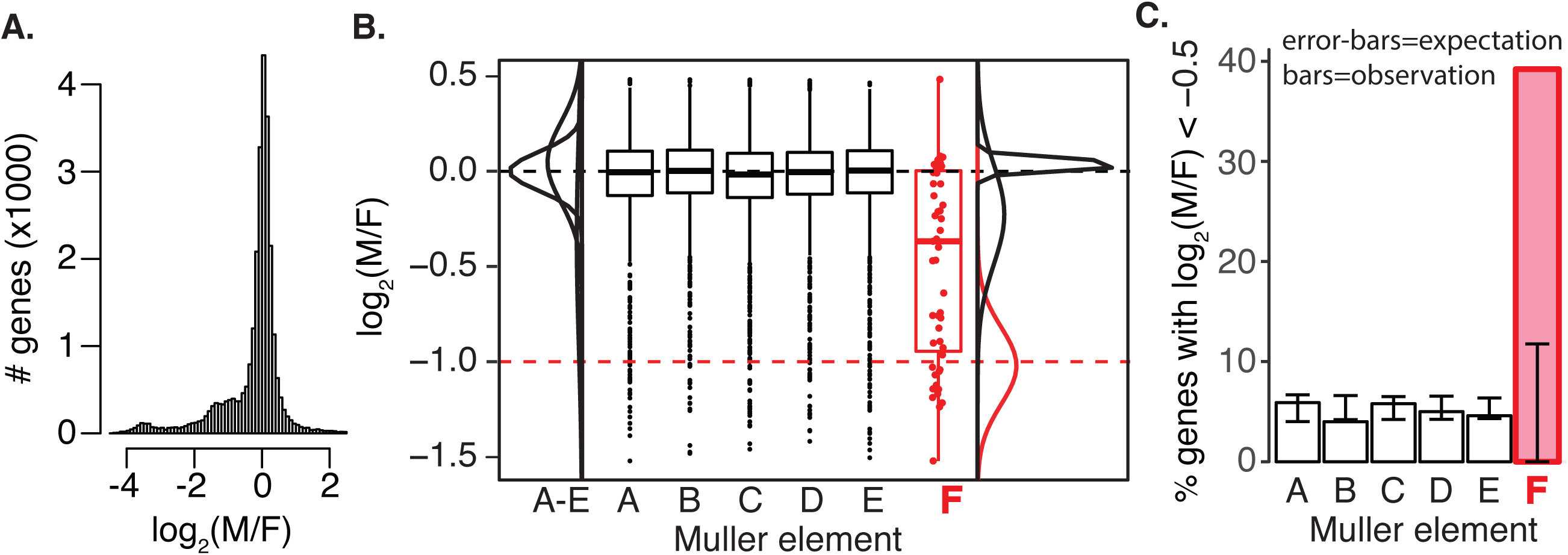
Reduced male-to-female sequence coverage of Muller element F homologs. **(A)** The distribution of 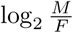 for all annotated genes in the *B. germanica* genome is shown, truncated to not include extreme values. **(B)** Boxplots show the distributions of 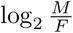 for *B. germanica* with homologs on one of the six *D. melanogaster* Muller elements. The red dashed line indicates the expectation of 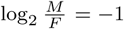 for X-linked genes. Each element F homolog is shown as a red dot on top of the box plot. The normal distributions from the mixture models for element A–E and element F homologs are shown next to the boxplots. **(C)** The percent of *B. germanica* genes with 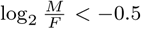 that have *D. melanogaster* homologs on each of the six Muller elements is plotted. The 95% confidence intervals (CIs) of the expected percent of genes for each Muller element are shown by the error bars. Observed percentages that lie outside the CI indicate an excess or deficiency of homologs on an element with moderately female-biased coverage.

To test the hypothesis that the German cockroach X chromosome is homologous to the ancestral brachyceran fly X (i.e., Muller element F), we evaluated if cockroach genes with *D. melanogaster* homologs on element F have lower 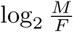 than genes with homologs on the other five elements. Cockroach genes with *D. melanogaster* homologs on Muller elements A–E have distributions of 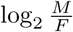 centered around 0, consistent with being autosomal (**Figure 2B**). In contrast, the 51 cockroach element F homologs have a median 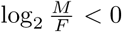, and the average 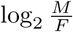 for element F homologs is significantly less than the other genes (*P* =10^*-*10^ using a Mann-Whitney *U* test comparing element F homologs with elements A–E). If all element F homologs were X-linked in cockroach, we would expect the median 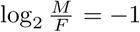 for genes with element F homologs. However, cockroach element F homologs have a median 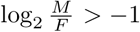. Therefore, we hypothesize that a disproportionate amount of, but not all, element F homologs are X-linked in German cockroach.

We next estimated the frequency of element F homologs that are X-linked in the German cockroach. First, we used the mclust package in R to fit a mixture of normal distributions to the 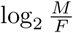 values of element F homologs [48]. The best fitting mixture consists of three distributions, with one centered at a mean of *-*1.02 (**Table 1**), close to the expectation of 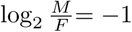 for X-linked genes. This suspected X-linked distribution contains ∼41% of the 51 element F homologs, and it has very little overlap with the other two distributions (**Figure 2B**). One of the other two distributions is centered very close to 0 (the expectation for autosomal genes), and it has very low variance. The third distribution has a mean 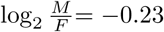 and a large variance. We suspect that the two distributions with 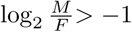 correspond to element F homologs that are autosomal in *B. germanica*. These two distributions may be the result of fitting normal distributions to a single non-normal distribution with a mode at 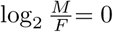 and a long tail extending into 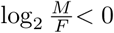. Consistent with this hypothesis, when we fit a mixture of two normal distributions to the 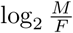 values of element F homologs, we obtain one distribution with a mean 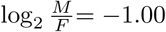 that has 43% of element F homologs, and a second distribution with a mean 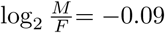 that has 57% of element F homologs (Additional file 2). Moreover, with a mixture of four normal distributions, we recover two distributions centered near 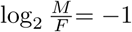 that together have 40% of element F homologs. Therefore, regardless of the number of distributions in our mixture model, we recover at least 40% of cockroach element F homologs that fall within a distribution consistent with X-linkage.

**Table 1.** Counts and proportions of genes assigned to each normal distribution in a mixture model of 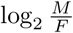 values. Genes have homologs in *D. melanogaster* on either element F or elements A–E.

| Distribution | proportion | mean | variance |
| --- | --- | --- | --- |
| *Element F genes* (51 total) |  |  |  |
| 1 | 0.41 | -1.020 | 0.041 |
| 2 | 0.36 | -0.225 | 0.069 |
| 3 | 0.23 | 0.038 | 0.001 |
| *Element A–E genes* (5,602 total) |  |  |  |
| 1 | 0.01 | -0.888 | 5.600 |
| 2 | 0.41 | 0.001 | 0.010 |
| 3 | 0.41 | 0.049 | 0.045 |
| 4 | 0.17 | -0.348 | 0.329 |

In contrast to element F, the 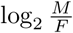 values for cockroach genes with *D. melanogaster* homologs on elements A–E can be best explained by a mixture of four distributions (**Table 1**). The distribution within this mixture model that is most consistent with X-linkage has a mean of *-*0.89, a large variance of 5.6, and contains only 37 of the 5,602 element A–E homologs. Most element A–E homologs (4,957) are assigned to two distributions with means of 0.0015 and 0.049, which are both consistent with autosomes (**Figure 2B**). Together, our analysis of mixture models suggest that a large fraction of element F homologs are X-linked in German cockroach, whereas the vast majority of element A–E homologs are autosomal.

The distributions of 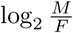 seem to describe two classes of element F homologs: autosomal genes with 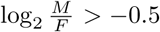, and X-linked genes with 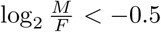 (**Figure 2B**). If there is an excess of element F homologs on the cockroach X, we expect a higher frequency of element F homologs to have 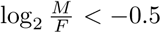 than genes on the other five elements. We therefore counted the number of genes with 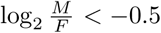 on each of the six Muller elements (**Table 2**). To determine a null distribution of those genes on each element, we randomly assigned the total number of genes with 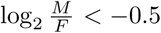 to the six elements based on the size of each Muller element (measured as the total number of cockroach genes on the element) in 1,000 bootstrap replicates of the data. A significant excess of cockroach element F homologs have 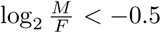 relative to our null expectation (**Figure 2C**). This provides further evidence that an excess of element F homologs is X-linked in German cockroach.

**Table 2.**
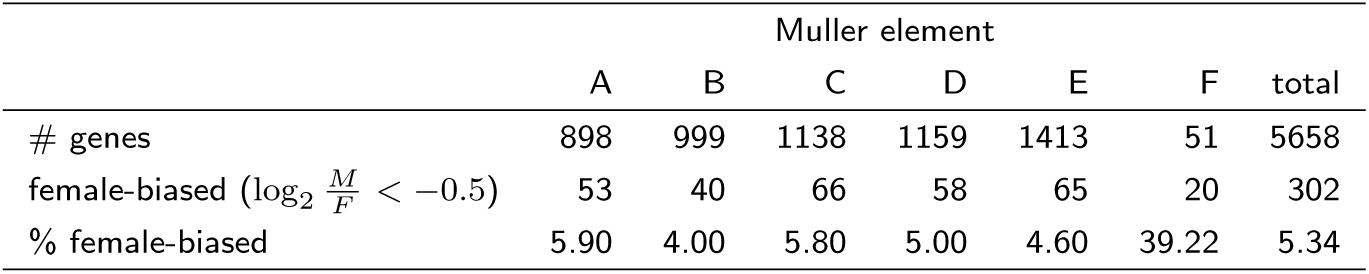
Genes with female-biased sequencing coverage and a *D. melanogaster* homolog on each Muller element.

|  | Muller element |  |  |  |  |  | total |
| --- | --- | --- | --- | --- | --- | --- | --- |
|  | A | B | C | D | E | F |  |
| # genes | 898 | 999 | 1138 | 1159 | 1413 | 51 | 5658 |
| female-biased ($\log_2 \frac{M}{F} < -0.5$) | 53 | 40 | 66 | 58 | 65 | 20 | 302 |
| % female-biased | 5.90 | 4.00 | 5.80 | 5.00 | 4.60 | 39.22 | 5.34 |

### Reduced heterozygosity of element F homologs in male cockroaches

German cockroach males have one copy of the X chromosome, and females have two copies of the X. We therefore expect that females could be heterozygous for polymorphic genetic variants in X-linked genes, whereas males must be hemizygous (only one allele per gene). If element F homologs are X-linked in cockroach, we expect to observe an excess of element F homologs without heterozygous variants in an individual male when compared to element A–E homologs and also when compared to female heterozygosity in element F homologs. To test this prediction, we used the available cockroach genome sequence data to identify heterozygous sequence variants in cockroach genes (Additional file 1).

The German cockroach genome project generated sequence data from a single male and single female of an inbred laboratory strain [47]. We therefore expect to observe no heterozygous variants in the male for X-linked genes, but the female could have heterozygous X-linked variants. However, there are also likely to be errors in variant calling and genotyping that could produce false positive heterozygous calls. Because of these false positives, we may observe heterozygous variants in element F homologs in the male even if the genes are X-linked. To address this limitation, we tested for reduced heterozygosity in element F homologs in the male, rather than an absence of heterozygous variants.

We first compared heterozygosity of cockroach genes in the male and female across Muller elements (**Figure 3**). In the female, there is not a significant difference in heterozygosity between genes assigned to element F and genes on the other five elements (*P* =0.32 in a Mann-Whitney *U* test). In contrast, male element F homologs have significantly fewer heterozygous variants than genes on elements A–E (*P* =0.017 in a Mann-Whitney *U* test). This reduced male heterozygosity in element F homologs is consistent with an excess of element F homologs on the German cockroach X chromosome.

**Figure 3.**
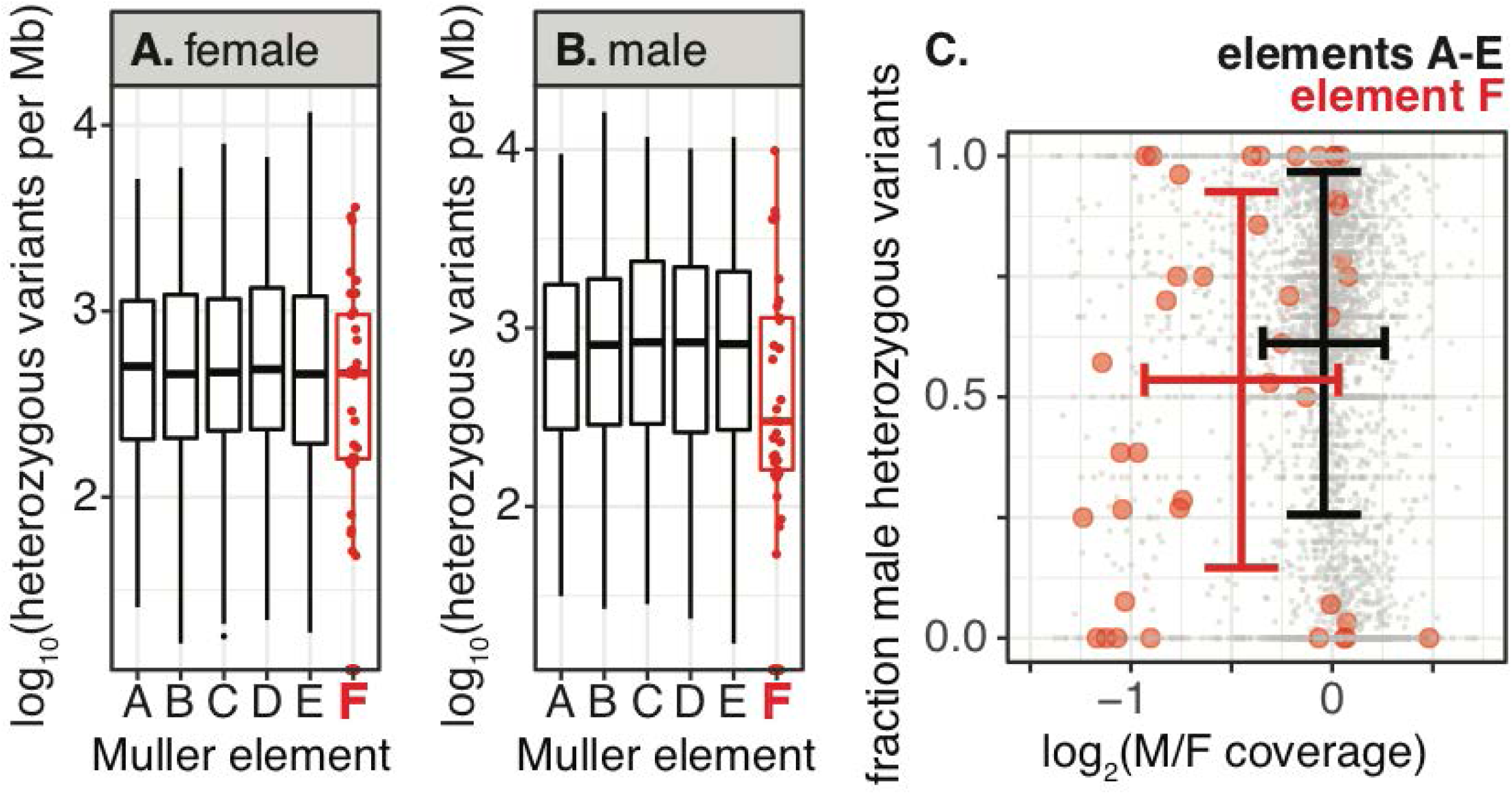
Reduced male heterozygosity in Muller element F homologs. **(A-B)** Boxplots show the distributions of heterozygous variants per Mb in males and females within genes assigned to each Muller element on a log_10_ scale. Each element F homolog is shown as a red dot on top of the box plot. **(C)** The points in the scatterplot show the 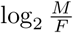 sequencing coverage and fraction of heterozygous variants in males for genes assigned to Muller elements, with element A–E homologs in gray and element F homologs in red. The standard deviations of 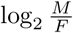 coverage and fraction of heterozygous variants in males are shown for element A–E homologs in black and element F homologs in red.

We expect candidate X-linked genes with reduced 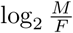 sequencing coverage to also have reduced heterozygosity in males relative to females. To test this hypothesis, we calculated, for each gene, a ratio of the number male heterozygous variants to the total number of heterozygous variants in the male and female samples. This value ranges from zero (if a gene only has heterozygous variants in the female) to one (if a gene only has heterozygous variants in the male). Equal heterozygosity in both sexes has a value of 0.5. Of the 40 element F homologs with sequencing coverage and heterozygosity data, 10 (25%) have both 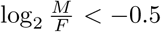 and fraction of male heterozygous variants *<* 0.5 (**Figure 3C**). This is significantly greater than the 2.5% of element A–E homologs with both 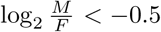 and fraction of male heterozygous variants *<* 0.5 (*z*=9.68, *P* =10^*-*21^). This result provides further evidence that there is an excess of element F homologs on the German cockroach X chromosome.

### Validation of candidate X-linked element F homologs

We selected two element F homologs that we hypothesize are X-linked (BGER000638 and BGER000663) to validate using quantitative PCR (qPCR). Both genes have 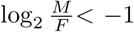, and one gene (BGER000638) has three times as many heterozygous variants in female compared to male (Additional file 1). The other gene has no heterozygous variants in either sex. We found that both genes had a significantly higher concentration in females relative to males in our qPCR assay, with an estimated female concentration that is twice the male concentration (Additional file 3). This is the expected result if both genes are X-linked. Therefore, male:female sequencing coverage, heterozygosity, and qPCR provide consistent evidence that element F homologs are X-linked in German cockroach.

### The cockroach X chromosome may be dosage compensated in males

We next tested if the haploid dosage of element F homologs affects their expression in cockroach males. The ideal data to test for the effects of a haploid X are expression measurements from males and females from the same tissue and developmental stage [10, 11]. Unfortunately, there are no available sex-matched RNA-seq gene expression datasets from German cockroach. We therefore used an alternative approach in which we compared expression in adult male heads with a mixed sex adult head sample (Additional file 1). We also compared expression in adult male heads with whole adult females (Additional file 1). If the haploid X chromosome is dosage compensated in males, we expect the distributions of log_2_ fold-change (log_2_ FC) expression between the two tissue samples to be equivalent for cockroach genes with homologs on element F and elements A–E. Indeed, there is not a significant difference in the median log_2_ FC between element F homologs and element A–E homologs (*P* =0.15 for male head *vs* mixed sex head, *P* =0.30 for male head *vs* whole adult female, with both *P*-values from Mann-Whitney *U* tests; **Figure 4A-B**).

**Figure 4.**
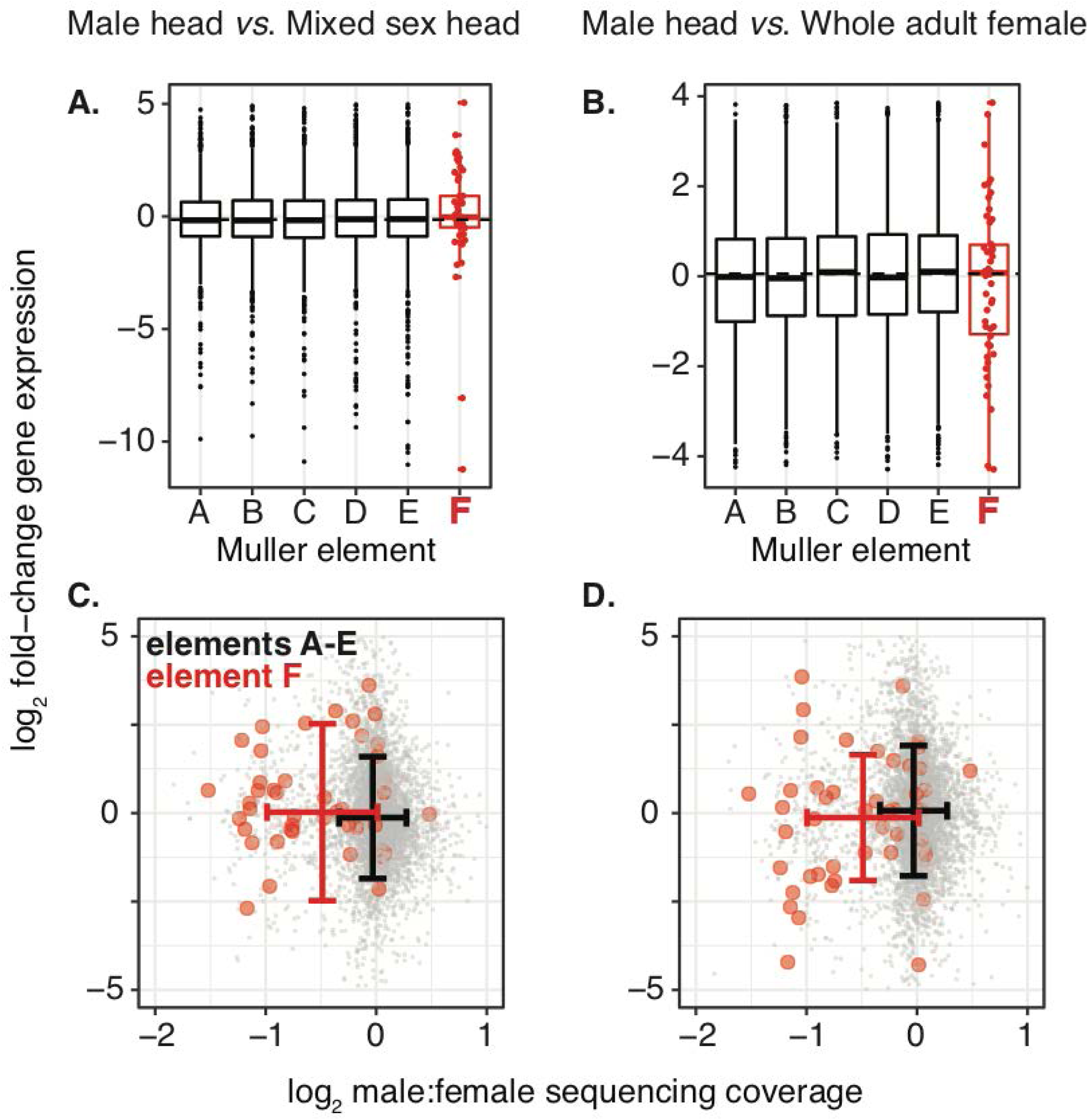
No reduced male expression of element F homologs. **(A-B)** Boxplots show the distributions of log_2_ FC of expression between either male and mixed sex heads or male heads and female whole adults for genes with *D. melanogaster* homologs on each Muller element. Each element F homolog is shown as a red dot on top of the box plot. **(C-D)** The points in the scatterplots show the 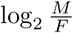 sequencing coverage and log_2_ FC of expression for genes assigned to Muller elements, with element A–E homologs in gray and element F homologs in red. The standard deviations of 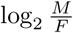 coverage and log_2_ FC expression are shown for element A–E homologs in black and element F homologs in red.

Only a subset of element F homologs is expected to be X-linked in cockroach based on 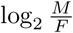 sequencing coverage (**Figure 2B**). If the X chromosome is dosage compensated in males, we expect the average log_2_ FC expression between tissue samples to be similar for element F homologs with evidence of X-linkage 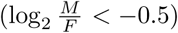 and element F homologs that appear to be autosomal 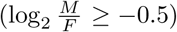. Indeed, there is not a significant difference in log_2_ FC between the two subsets of element F homologs (*P* =0.84 for male head *vs* mixed sex head, *P* =0.30 for male head *vs* whole adult female, with both *P*-values from Mann-Whitney *U* tests; **Figure 4C-D**). The same is true for element A–E homologs: there is not a significant difference in log_2_ FC of male head *vs* mixed sex head between low and high coverage element A–E homologs (*P* =0.054 in a Mann-Whitney *U* test), nor is there a significant difference in log_2_ FC of male head *vs* whole adult female between low and high coverage element A–E homologs (*P* =0.65 in a Mann-Whitney *U* test). The comparison of log_2_ FC in male *vs* mixed sex head for element A–E homologs has the lowest P-value. If this low P-value were evidence for a lack of dosage compensation, we would expect genes with low male sequencing coverage 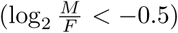 to have lower male expression than genes with higher male sequencing coverage 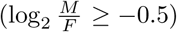 However, genes with low male sequencing coverage have higher male expression (median log_2_ FC= 0.0039) than genes with higher male sequencing coverage (median log_2_ FC= *-*0.15). Therefore, the limited RNA-seq data that are available suggest that the German cockroach X chromosome may be dosage compensated in males.

### Conservation of element F transcriptional regulators in cockroach

In some fly species where element F is the X chromosome, X-linked genes are present in a single (haploid) copy in males [27]. Males of the blow fly *L. cuprina* are haploid for such an X chromosome, and their X-linked genes are up-regulated by an RNA-binding protein encoded by a homolog of *Drosophila Pof* [41, 43]. POF localizes nearly exclusively to element F gene bodies in *D. melanogaster* [40, 49–51]. There is a *Pof* homolog in the cockroach genome (BGER016147), which we aligned to the *D. melanogaster* protein sequence. The most conserved region of *D. melanogaster Pof* overlaps with a predicted RNA binding domain within the cockroach protein sequence (**Figure 5A–B**). Therefore, a key component of the molecular machinery which regulates dosage compensation on the X-linked fly element F is present in the German cockroach genome.

**Figure 5.**
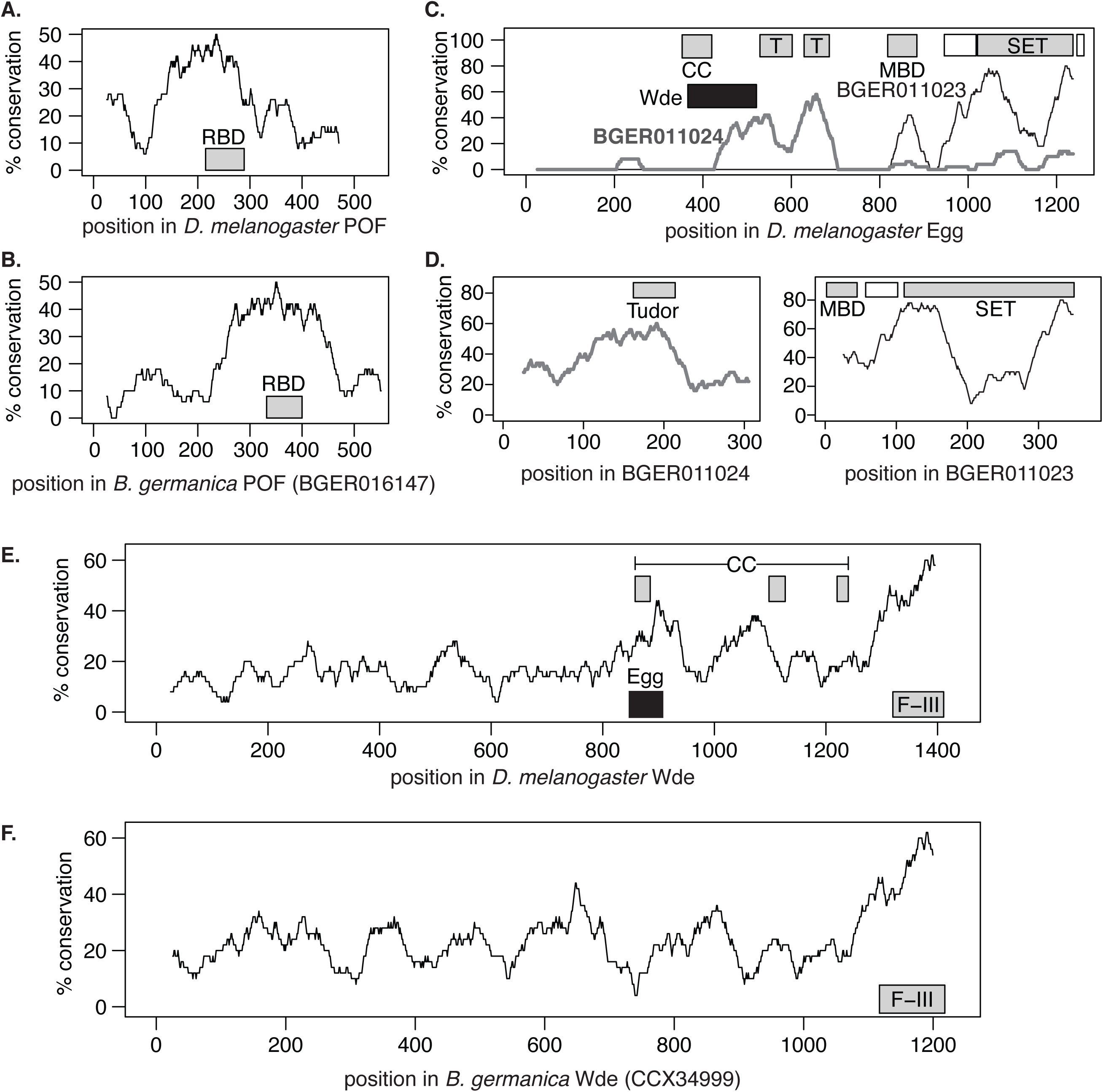
Three key regulators of element F transcription and chromatin are conserved in cockroach. Lines show the percent amino acid sequence conservation. The coordinates of the following predicted functional domains are shown as gray boxes in each graph: **(A-B)** RNA binding domain (RBD); **(C-D)** coiled-coil domain (CC), tudor domain (T), methyl-CpG-binding domain (MBD), and SET domain; **(E-F)** CC domain, and fibronectin type III repeats (F-III). **(C-D)** Predicted pre-SET domains are shown as white boxes next to SET domains. **(C)** The region of *D. melanogaster* Egg that interacts with Wde is shown by a black box, **(E)** as is the region of Wde that interacts with Egg.

The proteins encoded by *eggless* (*egg*) and *windei* (*wde*) interact with POF to create an environment around genes on element F that resembles pericentromeric heterochromatin in *Drosophila*. Egg is a SETDB1 homolog that is responsible for di- and/or tri-methylation of lysine 9 in histone H3 in the gene-dense region of *D. melanogaster* element F [52–56]. There are two predicted homologs of *egg* in the cockroach genome (BGER011023 and BGER011024). BGER011023 has a predicted SET lysine methyltransferase domain and a methyl-CpG binding domain commonly found in histone methyltransferases. BGER011024, on the other hand, has a tudor domain, which is found proximal to the SET domain in *D. melanogaster* Egg [57]. These predicted functional domains overlap with the portions of the cockroach proteins that are most conserved relative to *D. melanogaster* Egg (**Figure 5C–D**). BGER011023 and BGER011024 are contiguous on a single *B. germanica* scaffold (Scaffold202; KN196692), suggesting that together they may constitute a single gene that encodes all Egg functional regions.

Wde is an essential co-factor of Egg [58]. There is one predicted homolog of *wde* in the cockroach genome annotation (BGER025676), but an independently sequenced cockroach *wde* gene (CCX34999) is longer than the *wde* homolog predicted by the automated annotation [59]. We therefore compared CCX34999 with *D. melanogaster* Wde. CCX34999 contains a predicted fibronectin type-III domain at the C-terminal end, similar to *D. melanogaster* Wde [57]. The C-terminal end of CCX34999 is also the most conserved part of the protein relative to *D. melanogaster* Wde (**Figure 5E–F**). There is a coiled-coil region of *D. melanogaster* Wde that is required to interact with Egg. That coiled-coil region of Wde, and the corresponding region of Egg that interacts with Wde, are among the most conserved regions of the *D. melanogaster* proteins when compared to the cockroach homologs (**Figure 5C,E**). Therefore, homologs of *Pof* and its two key interactors are present in the German cockroach genome, showing it is possible that a similar mechanism may dosage compensate the cockroach and ancestral fly X chromosomes in hemizygous males.

## Discussion

We provide two lines of evidence that the X chromosome of the German cockroach, *B. germanica*, is homologous to Muller element F, which is X-linked in most flies. First, there is reduced sequencing coverage of nearly half of the Muller element F homologs in male cockroach, consistent with a haploid dose of the X chromosome in males (**Figure 2**). Second, there is decreased heterozygosity of element F homologs in male cockroach, including those with reduced male sequencing coverage (**Figure 3**). We therefore hypothesize that element F is an ancient X chromosome that was present in the most recent common ancestor (MRCA) of flies and cockroaches, and it has been conserved as an X chromosome in the German cockroach and many fly species. An alternative explanation for the excess of element F homologs on the cockroach X chromosome is that those genes independently became X-linked in both cockroaches and flies.

There are at least four lines of evidence that favor the hypothesis that element F is an ancient X chromosome retained since the MRCA of cockroaches and flies, as opposed to convergent recruitment of the same genes onto the fly and cockroach X. First, an independent analysis concluded that the MRCA of flies and cockroaches had XX females and either XY or XO males [16]. Second, the *B. germanica* X chromosome stains heavily for heterochromatin [46], similar to the brachyceran fly X-linked element F [60]. X chromosomes tend to be euchromatic in males [34–36], making the similarity between the *B. germanica* and brachyceran X heterochromatin notable. However, most of what we know about insect sex chromosome heterochromatin comes from cytological examination of meiotic cells from testes [61], where sex-chromosome-specific heterochromatization could differ from the normal behavior in somatic cells [62]. Additional work is necessary to investigate the chromatin state of insect sex chromosomes outside of the male germline. Third, the observed number of element F homologs with evidence for X-linkage in cockroach greatly exceeds the expectation if the X chromosomes of flies and cockroaches were independently derived (**Figure 2C**). Fourth, the fraction of element F homologs that appear to be X-linked in cockroach (*>*40%) is consistent with two separate estimates of the expected conservation of a shared X chromosome that was present in the MRCA of flies and cockroaches. We explain the two separate estimates of expected X chromosome conservation below.

The first estimate of expected conservation of an X-linked element F draws upon rates of gene relocation between Muller elements in *Drosophila*. If element F were the ancestral X chromosome of the MRCA of flies and cockroaches, we would expect some relocation of genes onto and off of element F as the lineages leading to cockroaches and flies diverged from their MRCA [63]. Based on the frequency of gene relocation between Muller elements in *Drosophila* [64] and the sizes of the elements in *D. melanogaster*, we expect 6.4 genes to have relocated off element F in the cockroach lineage and 1.3 genes to have relocated onto element F in the fly lineage (see Methods for calculations). There are up to 30 (60% of 51) *D. melanogaster* element F homologs that do not have evidence for X-linkage in cockroach (**Figure 2B**). Gene movement alone can thus explain 7–8 of these apparently autosomal element F homologs.

The second estimate of expected conservation of an X-linked element F extrapolates from the conservation of element F between *D. melanogaster* and the blow fly *L. cuprina*. In the *L. cuprina* genome, only 67.1% (49/73) of genes with *D. melanogaster* element F homologs are X-linked [43]. Assuming a linear relationship between divergence time [37, 65] and conservation of element F gene content, we would expect only 11.1% of cockroach genes with element F homologs to be X-linked:

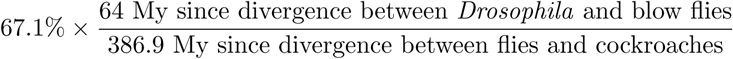

Our estimate of the fraction of element F homologs that are X-linked in *B. germanica* (*>*40%) is in between the estimates predicted based on rates of gene relocation and a linear loss of gene content. Therefore, conservation of an X-linked element F from the MRCA of flies and cockroaches is consistent with the expected amount of gene movement in the time since the MRCA.

Curiously, there is a long tail of genes with much higher sequencing coverage in females relative to males 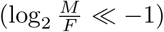, regardless of the Muller element of their *D. melanogaster* homologs (**Figure 2A**). Sexually dimorphic amplification (endoreplication) of a subset of the genome has been documented in insects, such as in the chorion genes that are highly expressed in the *Drosophila* ovary [66, 67]. It is therefore possible that a subset of the cockroach genome is disproportionately amplified in females (possibly to meet the gene expression demands of oogenesis), causing the long tail of negative 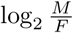 values that we observe. Additional work is necessary to test this hypothesis.

Our analysis of RNA-seq data suggests that the cockroach X chromosome may be dosage compensated in males—we find no evidence for reduced expression of element F homologs in male cockroaches, regardless of whether the genes appear to be haploid in males (**Figure 4**). Previous work found evidence that the cockroach *tra* homolog may regulate dosage compensation because knockdown of *tra* in cockroach females results in female-specific lethality of their progeny [68]. Here, we found that homologs of genes involved in regulating expression of element F genes in flies are present in the cockroach genome, with their functional domains conserved (**Figure 5**). This is consistent with cockroaches and flies sharing a mechanism of X chromosome dosage compensation that has been conserved since their MRCA. Future work should further investigate if the regulators of sex determination and dosage compensation in flies (e.g., *tra* and *Pof*) have similar roles in cockroach. An important limitation of our analysis is that we did not compare the same tissues between males and females [10, 11]. Our inference of dosage compensation may be confounded by, for example, differences in cell types between tissues [69]. Further work is therefore necessary to more rigorously test for dosage compensation of the cockroach X chromosome with appropriate gene expression comparisons between males and females.

Finally, our results provide evidence that X chromosomes can be conserved even though there are changes in the master regulators of sex determination. Sex in *B. germanica* is likely determined by X chromosome dosage, analogous to *Drosophila*, but different from the ancestral fly sex determination system, which relies on a dominant male determiner located on the Y chromosome (**Figure 1**). It is unlikely that the same X-linked dosage sensitive factors determine sex in cockroach and *Drosophila* because the X chromosome is not homologous between the two taxa (element A is the X chromosome in *Drosophila*). In addition, the master regulators of *Drosophila* sex determination almost certainly differ from the sex determiners in the MRCA of brachyceran flies, which likely used a Y-linked male-determiner (**Figure 1**). Moreover, the sexually dimorphic splicing of the sex determination pathway gene *tra* differs between German cockroach and flies [68]. Therefore, we hypothesize that *B. germanica* has a homologous X chromosome with the MRCA of brachyceran flies, but the sex determination system is not conserved between cockroaches and flies. Our results suggest that conservation of sex chromosomes does not necessarily imply conservation of sex determination. Future work addressing this problem could inform our understanding of how evolutionary transitions in sex determination pathways can be decoupled from sex chromosome turnover [70].

## Conclusions

We present evidence that the X chromosome of the German cockroach is homologous to an X chromosome shared by many fly species. We hypothesize that this X chromosome was inherited from the MRCA of cockroaches and flies *>*400 Mya. To the best of our knowledge, this would be the longest documented conservation of an X chromosome. This ancient X chromosome may be dosage compensated in male cockroaches and flies by a conserved mechanism. The extremely long-term conservation of the X chromosome is especially remarkable because cockroaches and flies have diverged in their sex determination pathways, suggesting that sex chromosome conservation can be decoupled from the evolution of sex determination.

## Methods

### Assigning German cockroach genes to Muller elements

*Drosophila* and other fly genomes are organized into six chromosomes (or chromosome arms) known as Muller elements [25, 38, 71, 72]. Muller element F is the ancestral X chromosome of brachyceran flies, and elements A–E are autosomal in flies with this ancestral karyotype [27]. We assigned each *B. germanica* gene with a single *D. melanogaster* homolog to the Muller elements of its homolog. We retrieved the *D. melanogaster* homologs of *B. germanica* genes from the Baylor College of Medicine i5k Maker annotation, version 0.5.3 [47]. This annotation pipeline was performed as part of the *B. germanica* genome project [47]. We only assigned *B. germanica* genes to Muller elements if they have a single *D. melanogaster* homolog in the annotation (i.e., we did not include genes with multiple predicted *D. melanogaster* homologs or without any predicted homologs).

### Differential sequencing coverage in males and females

We tested for genes that were sequenced at different depths in males and females as a way to identify X chromosome genes [27]. First, we aligned paired-end reads from three male cockroach whole genome sequencing libraries (SRX693111, SRX693112, and SRX693113) and one female library (SRX693110) to the reference *B. germanica* genome assembly [JPZV00000000.1; 47] using BWA-MEM with default parameters [73]. We then assigned mapped read pairs to genes (from the v. 0.5.3 i5k annotation) if the first (forward) read aligned to any portion of a gene sequence. We only considered the forward read because insert sizes differ across the available sequencing libraries, which could introduce biases in gene coverage if we allowed or required both forward and reverse reads to overlap genes. Considering only the forward read should decrease the effect of these biases because read lengths are the same (101 bp) across all libraries. We summed across libraries to determine the total number of reads mapped to each gene for each sex. We next divided the number of male-derived (female-derived) reads aligned to each gene by the total number of male-derived (female-derived) reads aligned to all genes to determine a normalized mapping coverage of male-derived (female-derived) reads for each gene (Additional file 1). We used these normalized counts to calculate the log_2_ male:female read mapping coverage 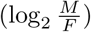 for each annotated cockroach gene, and we normalized the data so that the median across all genes assigned to Muller elements is zero.

We used the mclust package to fit a mixture of multiple normal distributions to the 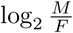 values [48]. We did this separately for element F homologs and genes assigned to elements A–E. The Mclust() function uses an expectation-maximization algorithm to obtain maximum likelihood estimators of the mean, variance, and number of genes in each normal distribution. It fits two different models for mixtures of 1 through 9 normal distributes: 1) mixture models where each normal distribution has the same variance (i.e., mixture of univariate normal distributions) and 2) mixture models where the normal distributions have unequal variances. We then compared Bayesian information criteria (BIC) across the nested models to determine the number of normal distributions that fit data the best (Additional file 2). We also compared BIC values to test if the best fitting distributions are univariate or have unequal variances.

### Quantitive PCR validation of candidate X-linked genes

We used qPCR to validate two candidate X-linked genes in German cockroach. Briefly, genomic DNA was extracted from the head and legs of five individual male and five individual female cockroaches from the Orlando Normal strain. We designed PCR primers to amplify the genomic region corresponding to each gene, as well as two control genes that we hypothesize are autosomal (sequences provided in Additional file 3). We used a StepOne Plus Real-Time PCR System (Applied Biosystems) to quantify the concentration of DNA from each of the candidate genes and the control genes in each individual cockroach. We then used a mixed effects model to assess the effect of sex on the concentration of the candidate X-linked genes. Details are provided in Additional file 3.

### Differential heterozygosity in males and females

We tested for genes with reduced heterozygosity in males (including relative to females) as an additional way to identify X chromosome genes. We used the Genome Analysis Toolkit (GATK) version 3.4-0 to identify heterozygous single nucleotide polymorphisms (SNPs) and small variants in the alignments of male and female sequencing reads described above, following the GATK best practices [74–76]. Because there is no reference variant set for cockroach, we used the following steps to extract high confidence variants [70]. First, we used Picard Tools version 1.133 to identify and remove duplicate reads, and we realigned indels with GATK. Then, we performed naïve variant calling using the GATK HaplotypeCaller with a phred-scaled confidence threshold of 20. We selected the highest confidence SNPs from that first-pass (QD < 2.0, MQ < 40, FS > 60, SOR > 4, MQRankSum < *-*12.5, ReadPosRankSum < *-*8). We also selected the highest confidence insertions and deletions (indels) from the first pass (QD < 2.0, FS > 200, SOR > 10, ReadPosRankSum < *-*20). We used those high quality variants to perform base recalibration, we re-input those recalibrated bases into another round of variant calling, and we extracted the highest quality variants. We repeated the process so that we had performed three rounds of recalibration, which was sufficient for convergence of variant calls. We applied GenotypeGVCFs to the variant calls from all of the Illumina libraries for joint genotyping of both males and females, and we selected only the high quality variants using VariantFiltration (FS > 30 and QD < 2). All three male sequencing libraries were treated as a single sample in this analysis because they came from the same individual male [47]. We used hard cut-off values because we did not have sufficient data to train a probabilistic variant filter. We then extracted variants that mapped to *B. germanica* genes (from the v. 0.5.3 i5k annotation). Variants were considered to be within a gene if they fell within the beginning and end coordinates of an annotated gene, including within exons or introns.

We identified heterozygous variants as those with two different alleles at that site in either the male or female sample. The two alleles could be either be one reference allele and one alternate, or they could be two alternate alleles. To calculate heterozygous variants per Mb within each gene, we used the differences of the beginning and end coordinates of each annotated gene in the genome assembly as a measure of gene length. To calculate the fraction of heterozygous variants in the male, we counted the number of heterozygous variants in the male (*H*_*m*_) and female (*H*_*f*_) samples separately for each gene. We then divided the number of heterozygous variants in the male sample by the sum of the number of heterozygous variants in the male and female samples for each gene (*H*_*m*_*/*[*H*_*m*_ + *H*_*f*_]).

### Differential gene expression using RNA-seq data

We compared the expression of genes in adult male heads (NCBI SRA accessions SRX3189901 and SRX3189902) with expression in a mixed sex adult head sample (SRX682022) using available RNA-seq data [77, 78]. We also compared male head expression with expression in whole adult females (SRX2746607 and SRX2746608) [47]. We aligned the RNA-seq reads from each library to *B. germanica* transcripts (from the version 0.5.3 i5k annotation) using kallisto [79]. The male head libraries were sequenced using single end reads, and we specified an average fragment length (-l) of 200 bp and a standard deviation (-s) of 20 bp. There is only a single transcript for each gene in the *B. germanica* annotation, and so we treated transcript-level read counts as equivalent to gene-wise counts. We also only included genes with at least 10 mapped reads across all samples. We then used DESeq2 to estimate the log2 fold-change of expression for each gene between male heads and mixed sex heads, as well as between male heads and whole adult females [80]. All reads from a given accession were treated as belonging to a single replicate (i.e., we summed read counts of different sequencing runs within each accession).

### Conservation of element F regulators

We aligned the sequences of three *D. melanogaster* proteins that regulate element F gene expression (POF, Eggless, and Windei) with their *B. germanica* homologs using MUSCLE [81]. We then calculated amino acid (aa) sequence conservation in 50 aa sliding windows (with 1 aa increments) in the reference protein sequence. Gaps in the cockroach sequences were counted as mismatches, and gaps in the *D. melanogaster* sequences were ignored. Functional domains were predicted by the NCBI Conserved Domain Database [57] or retrieved from UniProt [82].

### Expected conservation of element F

We performed calculations to estimate the number of genes relocated onto and off of element F in the lineages leading to cockroach and flies. First, the expected number of genes relocated from element F to the other elements in the lineage leading to the German cockroach was estimated from the observed number of X-to-autosome relocations in the lineage leading to *D. melanogaster* since the divergence with *Drosophila pseudoobscura* (24) [64], the fraction of genes on element F (86*/*14237 = 0.006) and element A (the *Drosophila* X chromosome, 2274*/*14237 = 0.16) in *D. melanogaster* [83], the divergence time between *D. melanogaster* and *D. pseudoobscura*(54.9 My) [84], and the divergence time between flies and cockroaches (386.9 My) [44]. We assumed that the rate of relocation from the ancestral X chromosome to the autosomes in the lineage leading to cockroach is the same as the rate from the *Drosophila* X to autosomes. We then calculated the expected number of genes relocated from element F to other elements in the lineage leading to the German cockroach as:

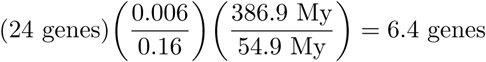

Second, to estimate the number of genes relocated onto element F from other elements in the lineage leading to *D. melanogaster*, we included an estimate of the number of autosome-to-X relocations in the lineage leading to *D. melanogaster* since the divergence with *D. pseudoobscura* (5) [64]. We treated element F as an X chromosome in the entire lineage leading from the MRCA of flies and cockroach, which it was for most of that time (332*/*387 My). We then calculated the expected number of genes relocated onto element F in the lineage leading to *D. melanogaster* as:

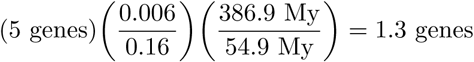

## Declarations

The data analyzed in the current study are available from the NCBI accessions provided in the Methods.

## Author’s contributions

RPM and JRW conceived the project. RPM analyzed the genomic data. JRW collected biological materials, extracted DNA, and designed primers for qPCR validation. PJD collected and analyzed the qPCR data. RPM wrote the paper, with assistance from PJD and JRW. All authors read and approved the final manuscript

## Acknowledgements

This work was supported by National Institutes of Health grant R35GM122592 to Artyom Kopp and National Institute of Food and Agriculture funds from the University of California. Computational analyses were performed on the Maxwell cluster provided by the Research Computing Data Core at the University of Houston.

## Additional Files

Additional file 1 — Gene coverage, heterozygosity, and expression

Data table showing read mapping coverage, heterozygosity, and gene expression analysis for each cockroach gene.

Additional file 2 — Mixture models fit to 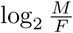 sequencing coverage

Table with properties of mixture models fit to 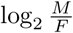 values for cockroach genes with *D. melanogaster* homologs on element F and elements A–E.

Additional file 3 — qPCR validation of 2 candidate X-linked genes

Methods and results of qPCR validation of cockroach genes with evidence of X-linkage.

