## Additional file 3 for "The X chromosome of the German cockroach, Blattella germanica, is homologous to a fly X chromosome despite 400 million years divergence"

### Quantitative PCR (qPCR) Validation of 2 Candidate X-linked Genes

We used qPCR to assess if the genes BGER000638 and BGER000663 are X-linked by testing if they are at twice the concentration in females as males relative to control genes. Both genes have a *D. melanogaster* ortholog on element F, and their male:female relative sequencing coverage ($log_2$M/F) is less than -1. In addition, BGER000638 has 3x as many heterozygous sites in the female sample (9 sites) than in the male sample (3 sites). We used putative single copy genes for RNA polymerase (*RNAPol*) and Triosephosphate isomerase (*tpi*) as internal reference autosomal genes (Wiegmann et al. 2009). We designed PCR primers to amplify BGER000638, BGER000663, *RNAPol*, and *tpi*, which we validated with the IDT OligoAnalyzer Tool (Table 1). The PCR primers were ordered from Sigma Life Science (Sigma-Aldrich, TX, USA).

| **Gene** | **Primer sequences** |
|---|---|
| *RNAPol* | Forward - 5'- GCGGCTGATGAGCAAACAGAGGC - 3'<br>Reverse - 5'- TGTTCAGTAGCTGCGACTGTAGCCAGAGT - 3' |
| *tpi* | Forward - 5'- CATTCGTAGGTTGTTCATAGCGTTCA - 3'<br>Reverse - 5' - GTTTCTGTCCGTTCTGTACCTCCATGTC - 3' |
| BGER000638 | Forward - 5'- GGTGATGCTGTACGCTATCTGCCTT - 3'<br>Reverse - 5'- GTTGGTCTCGTAACTCGTGAGCAAG - 3' |
| BGER000663 | Forward - 5'- AGAACGCCTTCACATGGTTGTACTTT - 3'<br>Reverse - 5'- CGAACTTCAACTGTGCTTCCTCCACGA - 3' |

**Table 1.** PCR primers used to test for X-linkage in qPCR assay.

We used a phenol:chloroform protocol to extract genomic DNA from five adult male and five adult female cockroaches from the Orlando Normal strain (obtained from Coby Schal's lab at North Carolina State University). DNA extractions were performed separately for each individual. Per individual, we collected tissue from all six legs and the head for DNA extraction. Tissue was digested for 3-4 hours with proteinase K (Sigma Aldrich P4850) at a concentration of 0.05 mg/mL, followed by a phenol:chlorofom DNA extraction. We performed two rounds of phenol:chloform mixing and centrifugation, followed by a round of mixing and centrifugation with chloroform alone. DNA was precipitated in ethanol and resuspended in Qiagen Elution Buffer.

The specificity of each primer pair was verified on three samples from each sex via PCR using the GoTaq Flexi DNA Polymerase kit. Each PCR contained 3 µl 5X Green GoTaq Flexi Buffer, 0.6 µl $MgCl_2$, 0.3 µl 2mM dNTP's, 0.6 µl each of 10 µM forward and reverse primers, 0.15 µl Flexi GoTaq DNA Polymerase, 9.25 µl nuclease-free $H_2O$ and 0.5 µl template genomic DNA. Amplifications were carried out in a Bio-Rad T100 thermal cycler. The reactions consisted of an initial denaturation step at 95 °C for 3 min, followed by 35 cycles of 95 °C for 30 sec, 60 °C for 30 sec and 72 °C for 30 sec. A final extension at 72 °C for 5 minutes was applied, followed by holding at 4 °C. PCR products were visualized by 1.5% agarose gel electrophoresis. All PCR primer pairs yielded a single product (i.e., one band), confirming the specificity of the primers.

Following PCR verification, we performed qPCR in a StepOne Plus Real-Time PCR System (Applied Biosystems). We first ran serial dilutions (5 concentrations diluted in a 1:5 ratio) of one of our male biological samples to generate a standard curve for each primer pair. Reactions were performed in MicroAmp 96-well plates (Applied Biosystems) in a final volume of 10 µl, following the manufacturer's protocol for standard cycling conditions. The slopes of the threshold cycle ($C_t$) against log of DNA concentration for a given primer pair were used to calculate amplification efficiency using the equation $E = [10^{(-1/\text{slope})} - 1] \times 100$, where $E$ denotes amplification efficiency as a percentage. Amplification efficiencies for the *RNAPol*, BGER000638, and BGER000663 primer pairs were all >90%, while *tpi* showed a lower efficiency of 77%. We therefore selected *RNAPol* as the internal reference for further analyses.

Next, we performed amplification reactions for each primer pair and biological sample (5 male and 5 female genomic DNA samples) in triplicate (i.e., three technical replicates per sample and primer pair). Since all technical replicates for each sample and primer pair could not fit in one 96-well plate, two technical replicates were conducted on one plate blocked within 4 rows and 11 columns each, and the third technical replicate was conducted on a second plate. The suitability of each primer pair was further verified by the inclusion of no-template controls and a final dissociation step run immediately following qPCR.

A $C_t$ value was obtained for each reaction, and concentrations for a given technical replicate were calculated using the slopes and intercepts obtained from the standard curve for a respective primer pair using the formula $\log_{10} X_i = S_i \times C_{ti} + B_i$ for gene $i$, where $X$ is concentration, and $S$ and $B$ are the slope and intercept generated from the standard curve, respectively. Relative experimental gene concentrations were then normalized by dividing the concentration for a given biological and technical replicate by the concentration observed for the matched batch technical replicate's internal reference gene. To determine the effect of sex on relative experimental gene concentrations, we applied a mixed-effects model of the following equation: $Y_i = \beta_0 + \beta X_i + u_i + v_i$ where $Y_i$ is the relative concentration of sample $i$ for a given gene of interest, $X$ represents the fixed effect of sex, and $u$ and $v$ are the random effects of biological and technical replicate, respectively. The model was run using the *nlme* package in R.

Linear mixed-effects models revealed significant effects of sex on the relative concentrations of both BGER000638 ($\beta$ = -18.58, $t_{14}$ = -6.16, $p < 1 \times 10^{-6}$) and BGER000663 ($\beta$ = -6.53 $t_{14}$ = -3.54, $p = 3.3 \times 10^{-3}$). Female concentrations were approximately twice as large as those of males (Figure 1), confirming that female *B. germanica* have twice the number of copies of both BGER000638 and BGER000663 as males. Therefore, our qPCR confirms that both BGER000638 and BGER000663 are X-linked.

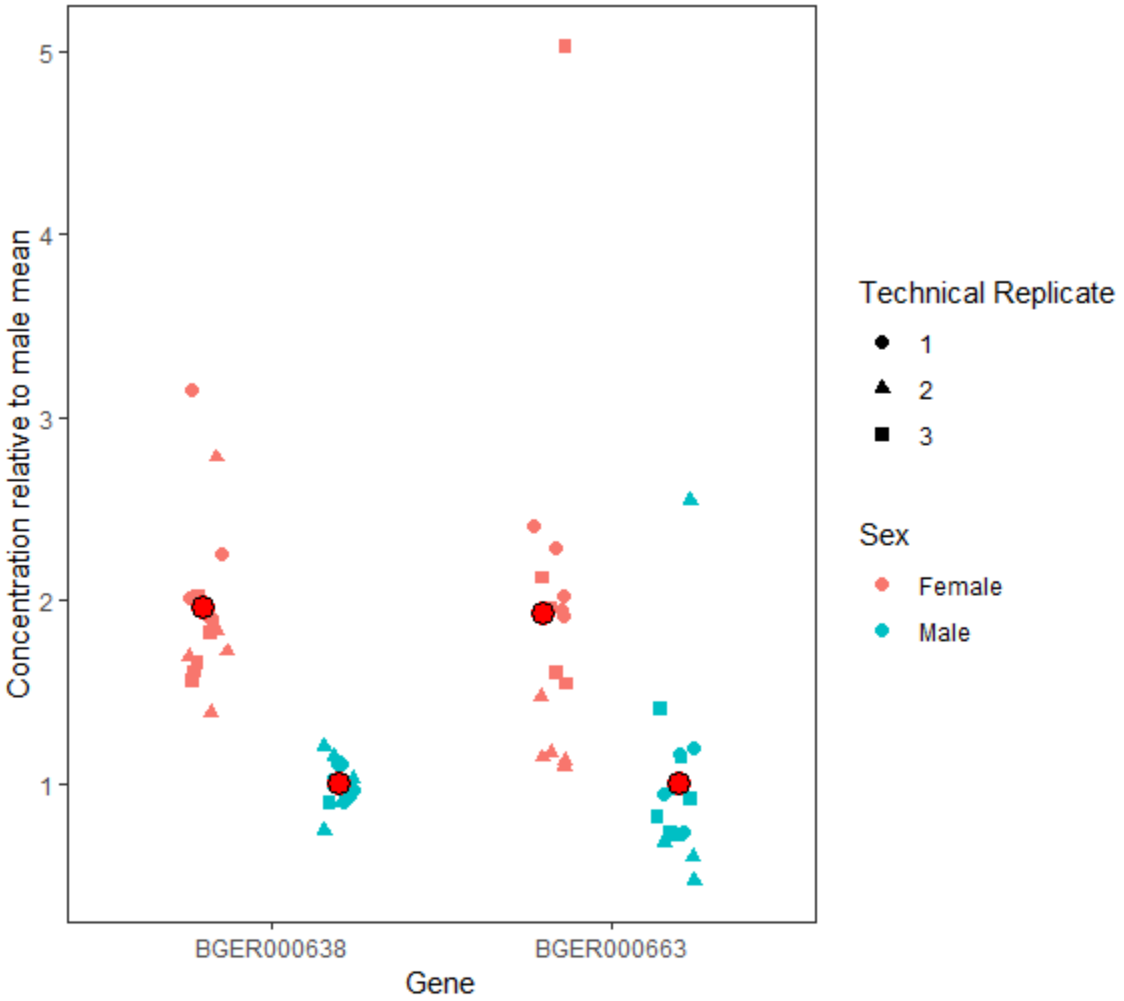


**Figure 1.** Relative concentrations of BGER000638 and BGER000663 in male and female *B. germanica*. Y-axis measures are relative to the mean male concentration for a given gene. Red circles denote within-sex means across all replicates.
